# Phase behaviour of C18-N-acyl sphingolipids, the prevalent species in human brain

**DOI:** 10.1101/2022.08.04.502749

**Authors:** Emilio J. González-Ramírez, Asier Etxaniz, Alicia Alonso, Félix M. Goñi

## Abstract

Lipidomic analysis of the N-acyl components of sphingolipids in different mammalian tissues had revealed that brain tissue differed from all the other samples in that SM contained mainly C18:0 and C24:1 N-acyl chains, and that the most abundant Cer species was C18:0. Only in the nervous system was C18:0 found in sizable proportions. The high levels of C18:0 and C16:0, respectively in brain and non-brain SM, were important because SM is by far the most abundant sphingolipid in the plasma membrane. In view of these observations, the present paper is devoted to a comparative study of the properties of C16:0 and C18:0 sphingolipids (SM and Cer) pure and in mixtures of increasing complexities, using differential scanning calorimetry, confocal microscopy of giant unilamellar vesicles, and correlative fluorescence microscopy and atomic force microscopy of supported lipid bilayers. Membrane rigidity was measured by force spectroscopy. It was found that in mixtures containing dioleoyl phosphatidylcholine, sphingomyelin and cholesterol, i.e. representing the lipids predominant in the outer monolayer of cell membranes, lateral inhomogeneities occurred, with the formation of rigid domains within a continuous fluid phase. Inclusion of saturated Cer in the system was always found to increase the rigidity of the segregated domains. C18:0-based sphingolipids exhibit hydrocarbon chain-length asymmetry, and some singularities observed with this N-acyl chain, e.g. complex calorimetric endotherms, could be attributed to this property. Moreover, C18:0-based sphingolipids, that are typical of the excitable cells, were less miscible with the fluid phase than their C16:0 counterparts. The results could be interpreted as suggesting that the predominance of C18:0 Cer in the nervous system would contribute to the tightness of its plasma membranes, thus facilitating maintenance of the ion gradients.

## INTRODUCTION

Sphingolipids, and in particular sphingomyelin (SM), the major sphingolipid in mammalian membranes, have long been understood as fundamentally structural components of the cell membrane. However, the discovery of the role of ceramide (Cer) as a signal of programmed cell death [1,2] changed that point of view, and focused the attention of molecular and cell biologists on the “simple” sphingolipids [3]. Cer and SM are related *structurally*, since SM is a Cer whose C1-OH has been substituted with phosphorylcholine, *metabolically*, since Cer is the product of SM hydrolysis by sphingomyelinase, and *functionally*, because SM is split into phosphorylcholine and Cer in response to a stress situation [4].

Studies from this and other laboratories, dating back from 1996 [5,6], have explored the behavior of Cer in phospholipid bilayers. Most of them have been reviewed in Castro et al. (2014), and in Alonso and Goñi (2018). One important property of sphingolipids in membranes is their capacity to form laterally segregated domains, enriched in SM and cholesterol (Chol), or SM and Cer. Formation of Cer-rich domains in fluid phosphatidylcholine (PC) bilayers was described by Huang et al. (1996). Collado et al. (2005) described the generation of domains in SM:Chol bilayers. Megha and London (2004) made the interesting observation that Cer displaced Chol from ordered lipid domains containing phosphatidylcholines, or SM, and Chol. Chiantia et al. (2006) observed coexistence of three kinds of domains in SM:PC:Chol:Cer supported lipid bilayers (SLB). Combining confocal fluorescence microscopy of giant unilamellar vesicles (GUV) with differential scanning calorimetry, Sot et al. (2008) described the Chol displacement by Cer in SM-containing liquid-ordered domains, and generation of gel regions in the giant vesicles. Later, Busto et al. (2010), using atomic force microscopy (AFM) showed how Chol displaced palmitoylceramide from its tight packing with palmitoylsphingomyelin in the absence of a liquid-disordered phase. Cholesterol interactions with ceramide and sphingomyelin were described by García-Arribas et al. (2016b), while the miscibility properties of Cer and SM were reviewed by Fanani and Maggio (2017). Al Sazzad et al. (2017) described how the long-chain sphingoid base of ceramides determined their propensity for lateral segregation. In the recent years, systems of increasing complexity, including glycerophospholipids, sphingolipids and Chol have been examined in this laboratory, integrating a variety of biophysical techniques, in particular DSC, confocal microscopy of GUV, and correlative AFM and fluorescence microscopy of SLB [17–20], as well as a detailed ^2^H-NMR study with selectively deuterated SM and Cer [21].

In a different series of experiments, Manni et al. (2018) performed an extensive scanning of the N-acyl components of sphingolipids in different mammalian tissues and cultured cells, using lipidomic technology. An unexpected result was that brain tissue differed from all the other samples in that SM contained mainly C18:0 and C24:1, and that the most abundant Cer species was C18:0. In all other samples, the predominant species of Cer were C24:0/C24:1, while C16:0 and C24:1 predominated in SM. Only in the nervous system was C18:0 found in sizable proportions. The high levels of C18:0 and C16:0, respectively in brain and non-brain SM, were important because SM is by far the most abundant sphingolipid in the plasma membrane [22]. In view of these observations, the present paper is devoted to a comparative study of the properties of C16:0 and C18:0 sphingolipids (SM and Cer) pure and in mixtures of increasing complexity, including some with C24:1 fatty acids, with the aim of exploring putative advantages of the C18:0 sphingolipids in the nervous system function.

## MATERIALS AND METHODS

### Chemicals

1,2-dioleoyl-sn-glycero-3-phosphocholine (DOPC, 850375), N-stearoyl-D-erythro-sphingosylphosphorylcholine (18:0 SM, sSM, 860586), N-nervonoyl-D-erythro-sphingosyl-phosphorylcholine (24:1 SM, nSM, 860593), N-stearoyl-D-erythro-sphingosine (18:0 Cer, sCer, 860518), N-nervonoyl-D-erythro-sphingosine (24:1 Cer, nCer, 860525), cholesterol (Chol, 700000), and the lipophilic fluorescent probe 1,2-dioleoyl-sn-glycero-3-phosphoethanolamine-N-(lissamine rhodamine B sulfonyl) (Rho-PE) were purchased from Avanti Polar Lipids (Alabaster, AL). Methanol and chloroform were from Fisher (Suwanee, GA). Buffer solution, unless otherwise stated, was 20 mM PIPES, 1 mM EDTA, 150 mM NaCl, pH 7.4. All salts and organic solvents were of analytical grade.

### Liposome preparation

All lipid mixtures are given as mole ratios. Lipid vesicle preparation consisted of mixing the desired lipids dissolved in chloroform/methanol (2:1, v/v) and drying the solvent under a stream of nitrogen. The removal of undesired organic solvent was ensured by keeping the lipid film under high vacuum for 90 min. Multilamellar vesicles (MLV) were formed by hydrating the lipid film with the buffer solution at 90 °C, helping the dispersion with a glass rod. The samples were finally sonicated for 10 min in a bath sonicator at the same temperature, to facilitate homogenization.

### Differential Scanning Calorimetry (DSC)

The measurements were performed in a VP-DSC high-sensitivity scanning microcalorimeter (MicroCal, Northampton, MA). MLV to a final concentration of 1 mM were prepared as described above with a slightly different hydration step: instead of adding the buffer solution in a single step, increasing amounts of the solution were added, helping the dispersion by stirring with a glass rod. Then the vesicles were homogenized by forcing the sample 50-100 times between two syringes through a narrow tube (0.5 mm internal diameter, 10 cm long) at a temperature above the transition temperature of the lipid mixture. Before loading the MLV sample into the appropriate cell both lipid and buffer solutions were degassed. 0.5 mL of suspension containing 1 mM total lipid concentration were loaded into the calorimeter, performing 8 heating scans at a 45 °C/h rate, between 10 and 90 °C for all samples. Phospholipid concentration was determined as lipid phosphorus and used together with data from the last scan to obtain normalized thermograms. After measurements, the software Origin 7.0 (MicroCal) provided with the microcalorimeter was used to subtract the baseline from the sample scan. Moreover, a cubic adjustment was performed to obtain a flat line before and after the transition temperature. Finally, the phospholipid concentration was determined using a phosphorus assay and the thermogram obtained was normalized with the concentration obtained. Finally, the different thermodynamic parameters from the scans were obtained. Further information on calorimetric studies of ceramide-containing samples can be retrieved elsewhere [23].

### Confocal microscopy of giant unilamellar vesicles (GUVs)

GUVs are prepared as described previously, using the electroformation method developed by Angelova and Dimitrov [24,25]. Lipid stock solutions were prepared in 2:1 (v/v) chloroform/methanol at a concentration of 0.2 mM, and appropriate volumes of each preparation were mixed. Labeling was carried out by pre-mixing the desired fluorescent probe (Rho-PE) with the lipids in organic solvent. Fluorescent probe concentration was 0.4 mol % Rho-PE. The samples were deposited onto the surface of platinum (Pt) wires attached to specially designed polytetrafluoroethylene (PTFE)-made cylindrical units that were placed under vacuum for 2 h to completely remove the organic solvent. The sample was covered to avoid light exposure. Then, the units were fitted into specific holes within a specially designed chamber (Industrias Técnicas ITC, Bilbao, Spain), to which a glass cover slip had been previously attached with epoxy glue. Once fitted, the platinum wires stayed in direct contact with the glass cover slip. The chamber was then equilibrated at the desired temperature by an incorporated water bath. 400 µL 10 mg/mL sucrose, prepared with high-purity water (SuperQ, Millipore, Billerica, MA) and heated at 90 °C, were added until the solution covered the Pt wires. The chambers were stopped with tightly fitting caps, and connected to a TG330 function generator (Thurlby Thandar Instruments, Huntingdon, UK). The alternating current field was applied with a frequency of 10 Hz and an amplitude of 940 mVrms for 120 min. The temperatures used for GUV formation were above the gel-to-liquid phase transition in all cases. The generator and the water bath were switched off, and vesicles were left to equilibrate at room temperature for 60 min. A slow cooling was needed to observe large enough domains by fluorescence microscopy [26]. After GUV formation, the chamber was placed onto a Leica epifluorescence microscope (DMI 4000B). The excitation wavelength was 561 nm. Emission was retrieved between 573 - 613 nm. Image treatment and quantitation were performed using the software Image J. No difference in domain size, formation, or distribution was detected in the GUV during the observation period or after laser exposure.

### Supported Planar Bilayer (SPB) formation

MLV were prepared as described above and introduced in a FB-15049 (Fisher Scientific Inc., Waltham, MA) bath sonicator, kept at 80 °C for 1 h. In this way, small unilamellar vesicles (SUV) were generated. SPB were prepared on high V-2 quality scratch-free mica substrates (Asheville-Schoonmaker Mica Co., Newport News, VA) previously attached to round 24 mm glass coverslips by the use of a two-component optical epoxy resin (EPO-TEK 301-2FL, Epoxy Technology Inc., Billerica, MA). SPB were obtained following the vesicle adsorption method [27,28]. Thereafter, 80 μL sonicated vesicles and 120 μL assay buffer containing 3 mM CaCl_2_ were added onto previously prepared 1.2 cm^2^ freshly cleaved mica substrate mounted onto a BioCell coverslip-based liquid cell for atomic force microscopy (AFM) measurements (JPK Instruments, Berlin, Germany). It has been described that the addition of divalent cations such as Ca^2+^ or Mg^2+^ enhances the vesicle adsorption process onto mica substrates [29]. Final lipid concentration was 150 μM. Vesicles were left to adsorb and extend for 30 min keeping the sample temperature at 60 °C. In order to avoid sample evaporation and ion concentration, the buffer was constantly exchanged with assay buffer without CaCl_2_ at 60 °C for the remaining time. Additional 30 min were left for the samples to equilibrate at room temperature, discarding the non-adsorbed vesicles by washing the samples 10 times with assay buffer without CaCl_2_, in order to remove remaining Ca^2+^ cations, which are reported to drastically affect the breakthrough force (F_b_) results of lipid bilayer nanoindentation processes [30]. The efficiency of repeated rinsing to obtain proper and clean supported lipid bilayers has been reported [31]. This extension and cleaning procedure allowed the formation of bilayers that did not cover the entire substrate surface. The presence of lipid-depleted areas helped with the quantification of bilayer thicknesses and the performance of proper controls for force-spectroscopy measurements. Planar bilayers were then left to equilibrate at room temperature for 1 h prior to measurements in order to avoid the presence of possible artifacts as segregated domains appear at high temperatures (over the T_m_) [32] and could still be present at lower temperatures if the cooling process was too fast (> 1 ºC/min) [29]. Finally, the BioCell was set to 23 °C to start the AFM measurements.

### Epifluorescence microscopy

Direct AFM-coupled inverted epifluorescence microscopy was performed in a Leica DMI 4000B microscope (Leica Microsystems, Wetzlar, Germany) using an appropriate filter cube for Rho-PE fluorochrome (excitation filter HQ545/30x, dichroic mirror Q570LP and emission filter HQ610/75m) (Chroma Tech., Bellows Falls, VT, USA). Images were acquired using a 100X/1.30 OIL objective (Leica Microsystems) with a high-resolution ORCA-R2 digital CCD camera (Hamamatsu Photonics, Shizuoka, Japan).

### AFM imaging

The topography of the supported planar bilayer was studied in an UltraSpeed AFM (JPK Instruments, Berlin, Germany) using the ‘QI Mode’, an imaging mode that performs force curves simultaneously at a low force. This ensures a lower tip-sample interaction than other methods available, which preserves the tip and the sample from damage. For proper measurements, the AFM was coupled to a Leica microscope and mounted onto an anti-vibration table and inside an acoustic enclosure (JPK Instruments). The BioCell liquid sample holder (JPK Instruments) was used in order to control the assay temperature at 23 °C. V-shaped MLCT Si3N4 cantilevers (Bruker, Billerica, MA) with nominal spring constants of 0.1 or 0.5 N/m were used for bilayer imaging, obtaining 256 × 256 pixel images though ‘QI Mode’. Images were line-fitted using the JPK Data Processing software.

### Force spectroscopy

V-shaped MLCT Si_3_N_4_ cantilevers (Bruker, Billerica, MA) with nominal spring constants of 0.1 or 0.5 N/m were individually calibrated in a lipid-free mica substrate in assay buffer using the thermal noise method. After proper bilayer area localization by AFM topography, force spectroscopy was performed at a speed of 1 µm/sec in no less than 500 × 500 nm bilayer areas in the form of 10×10 or 12×12 grids. Force steps were determined for each of the indentation curves as reproducible jumps within the extended traces. The resulting histograms were generated from at least 3 independent sample preparations with at least 3 independently calibrated cantilevers (n=300-1000). Control indentations were always performed in lipid-free areas before and after bilayer indentations to ascertain the formation of a single bilayer. The absence of artifacts or debris on the tip was assessed by the lack of any force-distance step on both trace and retrace curves.

## RESULTS

### 0. Study plan

The present work describes the phase behavior of 17 samples, pure lipids or lipid mixtures, mostly ternary or quaternary, in excess water (buffer), of which 1 contains nervonoyl (C24:1) sphingomyelin (nSM), 8 contain stearoyl (C18:0) sphingolipids (sSM and/or sCer) and the other 8 contain palmitoyl (C16:0) sphingolipids (pSM and/or pCer). All mixtures contain DOPC:SM:Chol at a 2:1:1 mol ratio; most of them contain Cer, at an additional 30 mol%, resulting in DOPC:SM:Chol:Cer 50:25:25:30 mol ratio mixtures, in which Cer is present at ≈23 mol%. Three molecular species have been tested for each sphingolipid, namely the C18:0, C16:0, and, occasionally, C24:1 acyl derivatives. All mixtures have been comparatively studied in couples, with C18:0 and C16:0 sphingolipids. They have been examined using DSC, confocal fluorescence microscopy of GUV, and AFM (topology and force spectroscopy) of supported lipid bilayers (SLB). In most cases, the SLB were doped with Rho-PE and their epifluorescence was recorded with a fluorescence microscope set up in corelation with the AFM microscope.

### 1. Pure C16:0 and C18:0 SM

As a preliminary part of this study, the DSC thermograms corresponding to pure C16:0 SM and C18:0 SM (pSM and sSM respectively) were recorded. The calorimetric traces are seen in Fig. 1A, B (curves I) and the thermodynamic data are collected in Table 1 (#1 and 2). The T_m_ of the main observed transitions were at 41.2 ºC and 44.1 ºC for the pSM and the sSM respectively, following the usual parallel increase in acyl chain length and T_m_ of saturated phospholipids [33]. In addition, a minor, rather wide peak at 30.2 ºC (Table 1) was detected for the pSM. The significance of this signal is unclear, but it might correspond to a gel-to-ripple phase transition that was observed with X-ray techniques [34]. Its area was ≈3% of the area of the main peak, and it was ignored in further discussions. It might be related to a larger peak observed with sSM, at 32.8 ºC. The latter was studied in detail by [35] and it was tentatively related to asymmetry between the sphingosyl and N-acyl chains in this and longer SM species. According to our previous study, the pSM molecule would be barely symmetric from the chain-length point of view.

**Figure 1.**
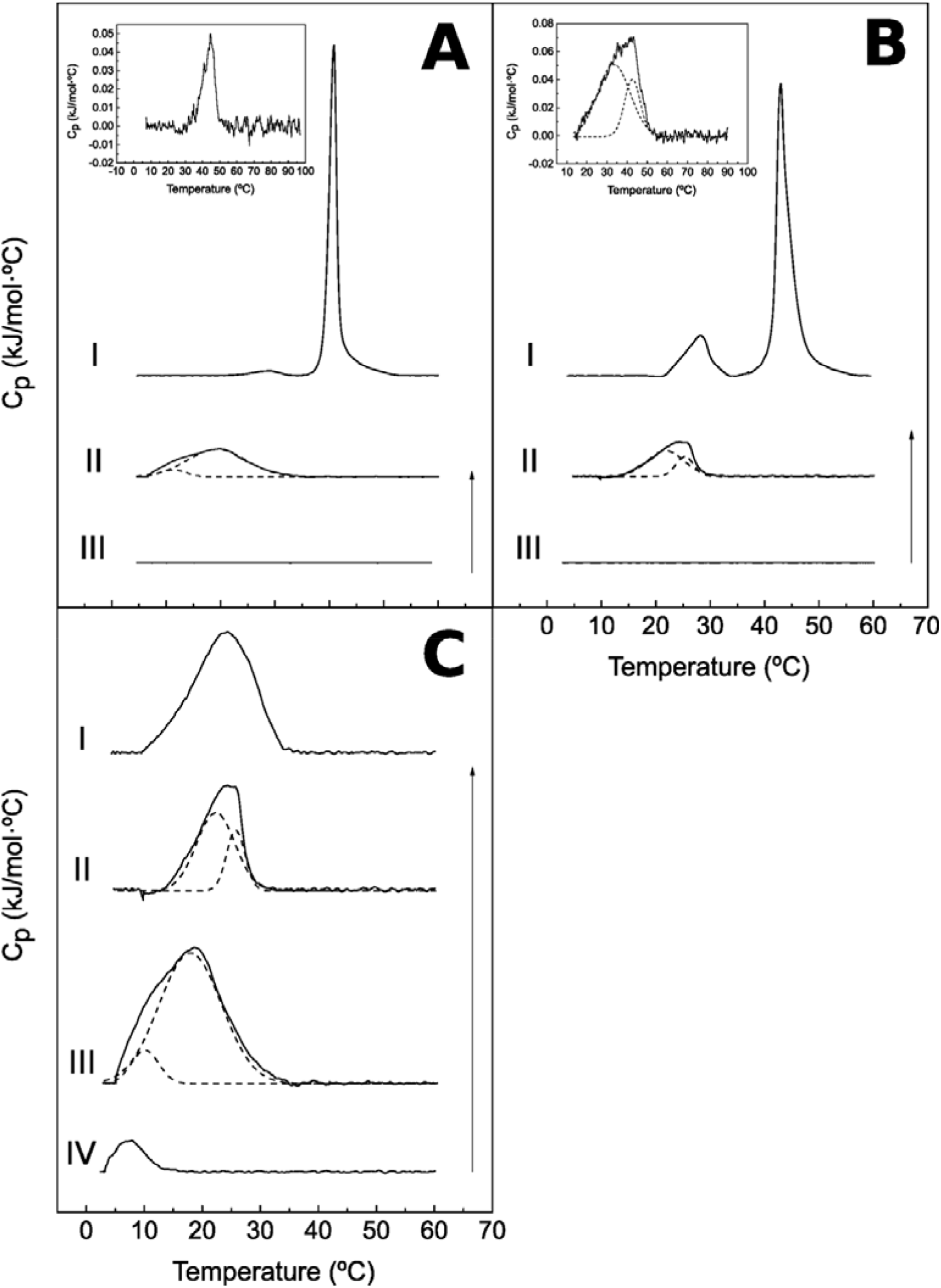
Representative DSC thermograms. **Panel A**: pSM (I), DOPC:pSM (2:1), and fitting with component curves (II), DOPC:pSM:Chol (2:1:1) (III), inset: zoom of thermogram III. **Panel B:** sSM (I), DOPC:pSM (2:1), and fitting with component curves (II), DOPC:sSM:Chol (2:1:1) (III), inset: zoom of thermogram III, and fitting with component curves. **Panel C:** DOPC:24:0 SM (2:1) (I), DOPC:sSM (2:1) (II), DOPC:pSM (2:1) (III), fitting with component curves in II and III, DOPC:nSM (IV). Thermograms C II and C III are taken, respectively, from B II and A II. Arrows: 5 kJ/mol·ºC.

**Table 1.**
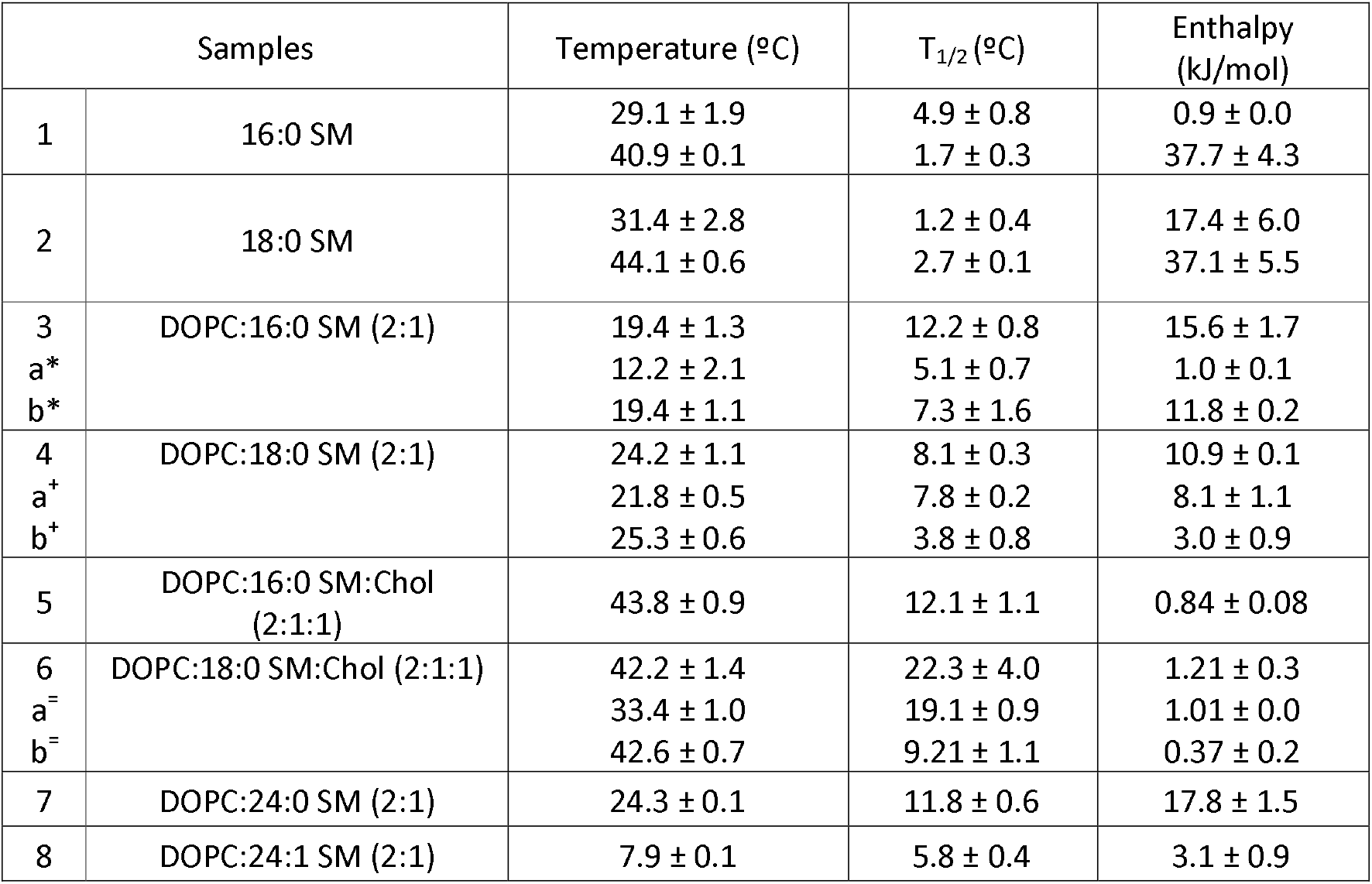
Thermodynamic parameters of the gel-fluid transitions, as obtained from DSC experiments. When a thermogram was fitted to various components, both the global and the component values are given, the latter denoted as a,b. Average values ± SD (n = 3). *Fitting R^2^ = 0.99. ^+^Fitting R^2^ = 0.99. ^=^Fitting R^2^ = 0.99.

| Samples |  | Temperature (°C) | T<sub>1/2</sub> (°C) | Enthalpy (kJ/mol) |
| --- | --- | --- | --- | --- |
| 1 | 16:0 SM | 29.1 ± 1.9 | 4.9 ± 0.8 | 0.9 ± 0.0 |
|  |  | 40.9 ± 0.1 | 1.7 ± 0.3 | 37.7 ± 4.3 |
| 2 | 18:0 SM | 31.4 ± 2.8 | 1.2 ± 0.4 | 17.4 ± 6.0 |
|  |  | 44.1 ± 0.6 | 2.7 ± 0.1 | 37.1 ± 5.5 |
| 3<br>a*<br>b* | DOPC:16:0 SM (2:1) | 19.4 ± 1.3 | 12.2 ± 0.8 | 15.6 ± 1.7 |
|  |  | 12.2 ± 2.1 | 5.1 ± 0.7 | 1.0 ± 0.1 |
|  |  | 19.4 ± 1.1 | 7.3 ± 1.6 | 11.8 ± 0.2 |
| 4<br>a+<br>b+ | DOPC:18:0 SM (2:1) | 24.2 ± 1.1 | 8.1 ± 0.3 | 10.9 ± 0.1 |
|  |  | 21.8 ± 0.5 | 7.8 ± 0.2 | 8.1 ± 1.1 |
|  |  | 25.3 ± 0.6 | 3.8 ± 0.8 | 3.0 ± 0.9 |
| 5 | DOPC:16:0 SM:Chol (2:1:1) | 43.8 ± 0.9 | 12.1 ± 1.1 | 0.84 ± 0.08 |
| 6<br>a=<br>b= | DOPC:18:0 SM:Chol (2:1:1) | 42.2 ± 1.4 | 22.3 ± 4.0 | 1.21 ± 0.3 |
|  |  | 33.4 ± 1.0 | 19.1 ± 0.9 | 1.01 ± 0.0 |
|  |  | 42.6 ± 0.7 | 9.21 ± 1.1 | 0.37 ± 0.2 |
| 7 | DOPC:24:0 SM (2:1) | 24.3 ± 0.1 | 11.8 ± 0.6 | 17.8 ± 1.5 |
| 8 | DOPC:24:1 SM (2:1) | 7.9 ± 0.1 | 5.8 ± 0.4 | 3.1 ± 0.9 |

### 2. DOPC:SM (2:1) binary mixtures

As a further step in this investigation, the calorimetric behavior of four DOPC:SM (2:1) binary mixtures was recorded. Representative DSC thermograms are shown in Fig. 1C, and the thermodynamic parameters are summarized in Table 1 (#3, 4, 7, 8). Pure DOPC does not exhibit any thermotropic phase transition in this temperature range (not shown). The gel-to-fluid T_m_ transition temperature of the binary mixtures decreases with the T_m_ of the corresponding pure SM, in the order C24:0 ≈ C18:0 > C16:0 > C24:1. Transition widths increase considerably when SMs are mixed with the unsaturated DOPC, as shown previously for egg SM [36]. The mixtures containing the unsaturated nSM are characterized by a relatively narrow thermogram (low T_1/2_) and, as expected from an unsaturated lipid, by a low transition enthalpy. The thermograms from the C18:0 and C16:0 mixtures were clearly asymmetric. Decomposing the overall thermogram into two components provided a good fit (R^2^ > 0.99 in both cases). In the C16:0-based mixture the main component melted at a higher temperature than the minor component (19.4 vs. 12.2 ºC), and the opposite was true for the components in the C18:0-based bilayers (respectively 21.8 and 25.3 ºC). These mixtures appeared homogeneous under the AFM and fluorescence microscopes (not shown), thus the presence of several components in the thermograms is probably due to nanodomains, on whose nature any comment would be mere speculation.

### 3. DOPC:SM:Chol (2:1:1) mixtures

#### 3.1. 16:0 (pSM) and 18:0 (sSM)

Representative DSC thermograms for these samples, namely DOPC:pSM:Chol (2:1:1) and DOPC:sSM:Chol (2:1:1), are shown in Fig. 1A, B. Mixing the saturated SM with unsaturated DOPC widens the transition of the former, as mentioned above (section 2). Moreover, adding Chol further spreads the endotherm, as first shown by D. Chapman and coworkers for DPPC [37], and later, for SM-containing mixtures, by Contreras et al. (2004). The gradual widening of the SM transition upon addition of DOPC and Chol is seen clearly when curves I, II and III are compared, respectively in Fig. 1A for pSM and in Fig. 1B for sSM. In the presence of both DOPC and Chol, a resetting of the Y-scale is necessary in order to visualize the transitions (insets in Fig. 1A, B).

The properties of these ternary mixtures are also summarized in Table 1 (#5, 6). Two marked differences are observed between the samples. One is that the calorimetric transition is much wider (higher T_1/2_) and asymmetric for the sSM-containing mixture. This may be related to the unusual fact of pure sSM exhibiting a 2-step gel-fluid transition, at 32.8ºC and 44.1ºC, when in pure form Jiménez-Rojo et al. (2014). It is conceivable that, in mixtures with other lipids, the 2-step transition will become a single, wider one.

The second important difference is detected examining the data in Fig. 2 and Table 2. While both mixtures give rise to lateral phase separation in GUV, the two phases exhibit rather different penetration forces with C16:0 (5.7 nN for the continuous phase and 11.3 nN for the segregated phase), but a single value of 2.7 nN with C18:0 (Table 2). In the latter case, the two phases differ in their heights, and presumably in composition, and yet they share the same penetration force. A similar circumstance has been described for the related mixture DOPC:lSM:nSM:Chol (2:0.5:0.5:1) in which two separated phases exhibited a single penetration force of 9.1 ± 1.4 nN, see Fig. 1 in González-Ramírez et al. (2020). The boundaries of the segregated domains are smooth, suggesting the coexistence of two fluid phases in these samples.

**Figure 2.**
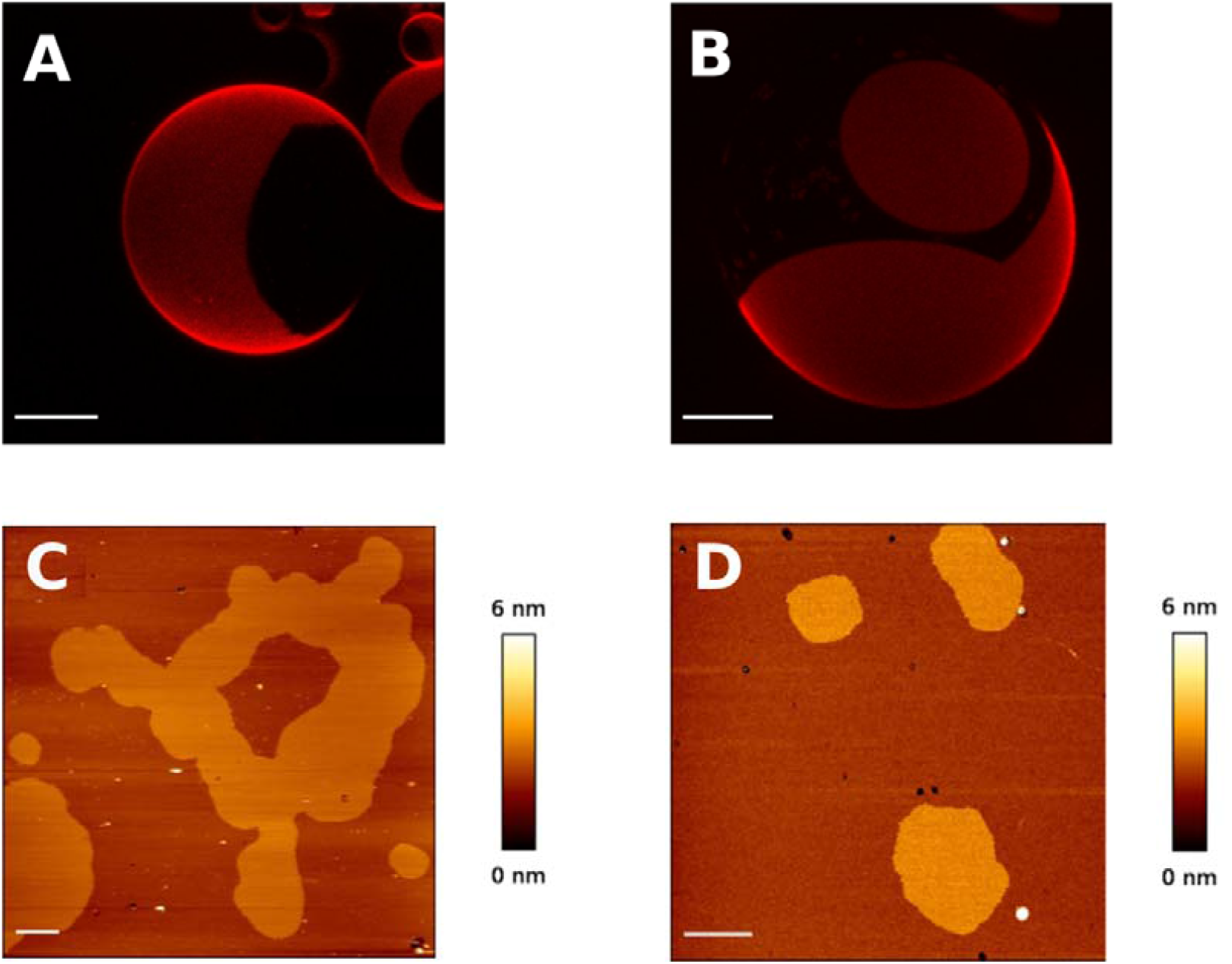
Giant unilamellar vesicles examined by fluorescence confocal microscopy and SLB topography images obtained with atomic force microscopy: DOPC:pSM:Chol (2:1:1) (A, C), DOPC:sSM:Chol (2:1:1) (B, D). Scale bar is 10 µm (GUV) and 1 µm (AFM).

**Table 2.** Physical characterization of DOPC:xSM:Chol (2:1:1) mixtures, with either pSM or sSM. AFM force spectroscopy: data from at least 3 independent sample preparations with at least 3 independently calibrated cantilevers. Average values ± S.D. (n=300-1000).

| Sample | x = 16:0 | x = 18:0 |  |
| --- | --- | --- | --- |
| Phases detected in GUV | 2 | 2 |  |
| AFM breakthrough forces (nN) | disc. $11.3 \pm 2.8$<br>cont. $5.7 \pm 1.3$ | $2.7 \pm 1.1$ | *** |

The observation of phase separation in both samples, both by AFM and by fluorescence microscopy (Fig. 2A - D), is related to the fact that the microscopic observations are performed at 23 ºC. At this temperature the thermotropic transition has probably started (considering the uncertainties of transition temperatures in GUV and SLB), thus phase coexistence is expected to occur [39].

### 2.2. (16:0 + 24:1) and (18:0 + 24:1) SM

These samples do not undergo gel-fluid calorimetric transitions in the T interval under study (4ºC-95ºC). GUV and topographic AFM images are shown in Fig. 3. DOPC:pSM:nSM:Chol (2:0.5:0.5:1) was studied by Maté et al. (2014), and it was found to be homogeneous in GUV according to fluorescence microscopy, and in SLB examined by AFM. Those authors associated the double bond in nSM to the lack of lateral heterogeneity in Chol-containing membranes. However the corresponding C18:0 sample, i.e. DOPC:sSM:nSM:Chol (2:0.5:0.5:1), did show phase separation, but with both phases sharing a single breakthrough force of 4.8 ± 0.7 nN. Thus the latter sample behaves in a way similar to the C18:0 mixture in Table 2. The domain boundaries in Fig. 3E, F are again rather smooth, an indication of fluid-fluid coexistence in the system. The presence of nSM tends to hinder phase separation and to fluidify the bilayers Maté et al. (2014). This is one case in which the behavior of a C18:0 sample departs from that of a C16:0 one. The formation of visible domains may be facilitated, or, conversely, mixing may be hindered, by the longer N-acyl chain.

**Figure 3.**
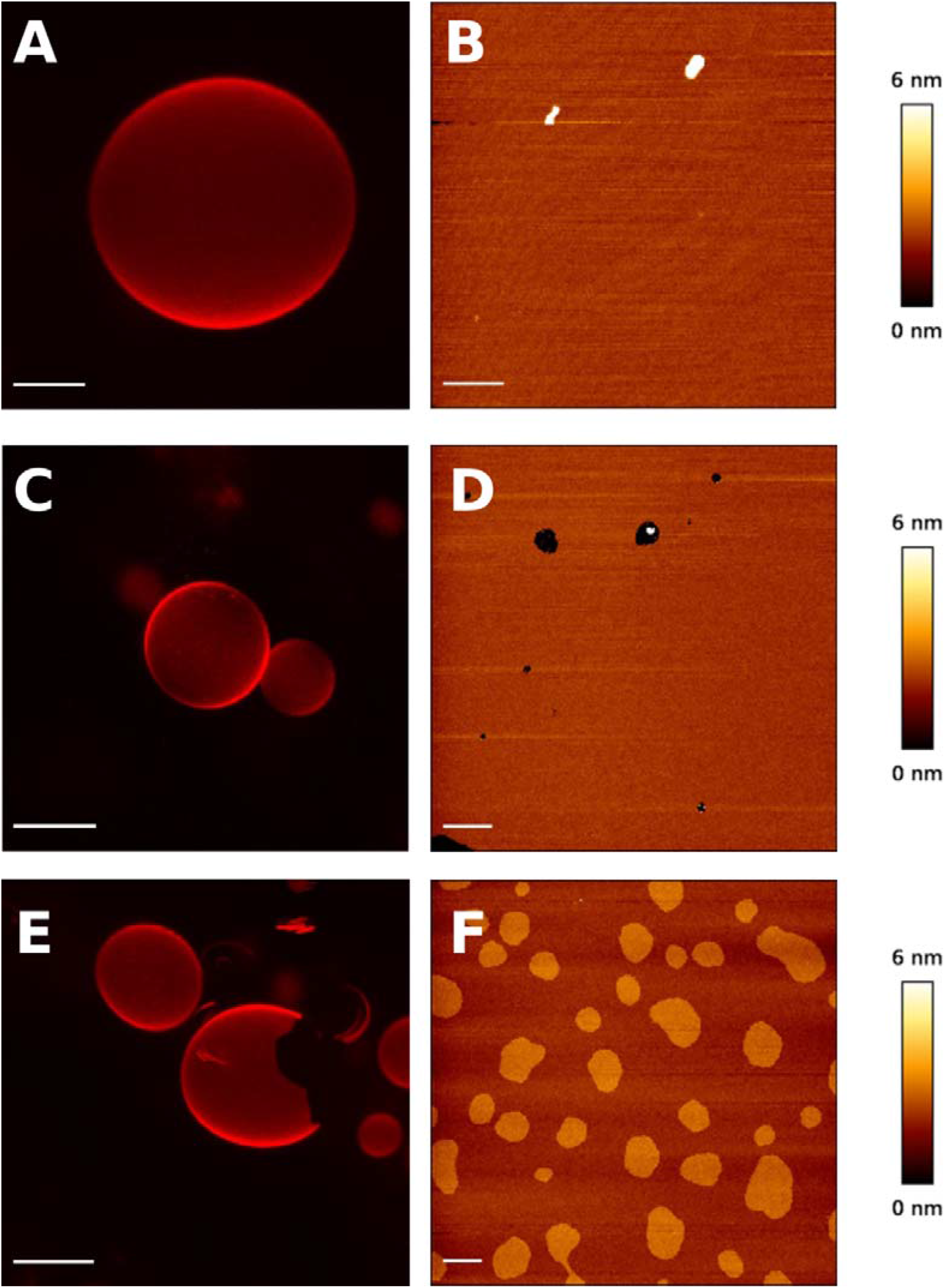
Giant unilamellar vesicles and bilayer topography images, obtained respectively with confocal fluorescence microscopy and with atomic force microscopy: DOPC:nSM:Chol (2:1:1) (A,B), DOPC:nSM:pSM:Chol (2:0.5:0.5:1) (C,D) and DOPC:nSM:sSM:Chol (2:0.5:0.5:1) (E,F). Scale bar is 10 µm (GUV) and 1 µm (AFM).

### 4. Mixtures containing DOPC:SM:Chol (2:1:1) + Cer

#### 4.1. DOPC:xSM:Chol (2:1:1) + 30 mol% saturated Cer

In these experiments, SM and Cer may contain either C16:0 or C18:0 acyl chains, the same chain in both lipids. The mol ratios in the mixture are DOPC:SM:Chol:Cer (50:25:25:30). The physical data are shown in Fig. 4 and in Table 4. From the calorimetric point of view, both samples exhibit a much larger transition enthalpy than those not containing Cer (compare Tables 1 and 4). This may be an indication that the observed endotherm arises from highly stable SM:Cer complexes. This behavior is observed as a rule in all our Cer-containing samples (see below), and it has been repeatedly observed in our previous studies [13,19,20,41]. Comparison of the C16:0 and C18:0 samples does not reveal major differences (even if the small variations observed can be statistically significant), except in the much increased transition enthalpy of the C18:0 mixture. Examination of the thermograms (Fig. 4A) suggests that, as seen in Fig. 1 for binary and ternary mixtures, the asymmetric sSM and sCer are giving rise to domains that can be identified calorimetrically. At least three components are required to fit the complex thermogram of the DOPC:sSM:Chol:sCer mixture (Fig. 4A, bottom). The one melting at a higher T corresponds probably to domains containing higher Cer proportions.

**Figure 4.**
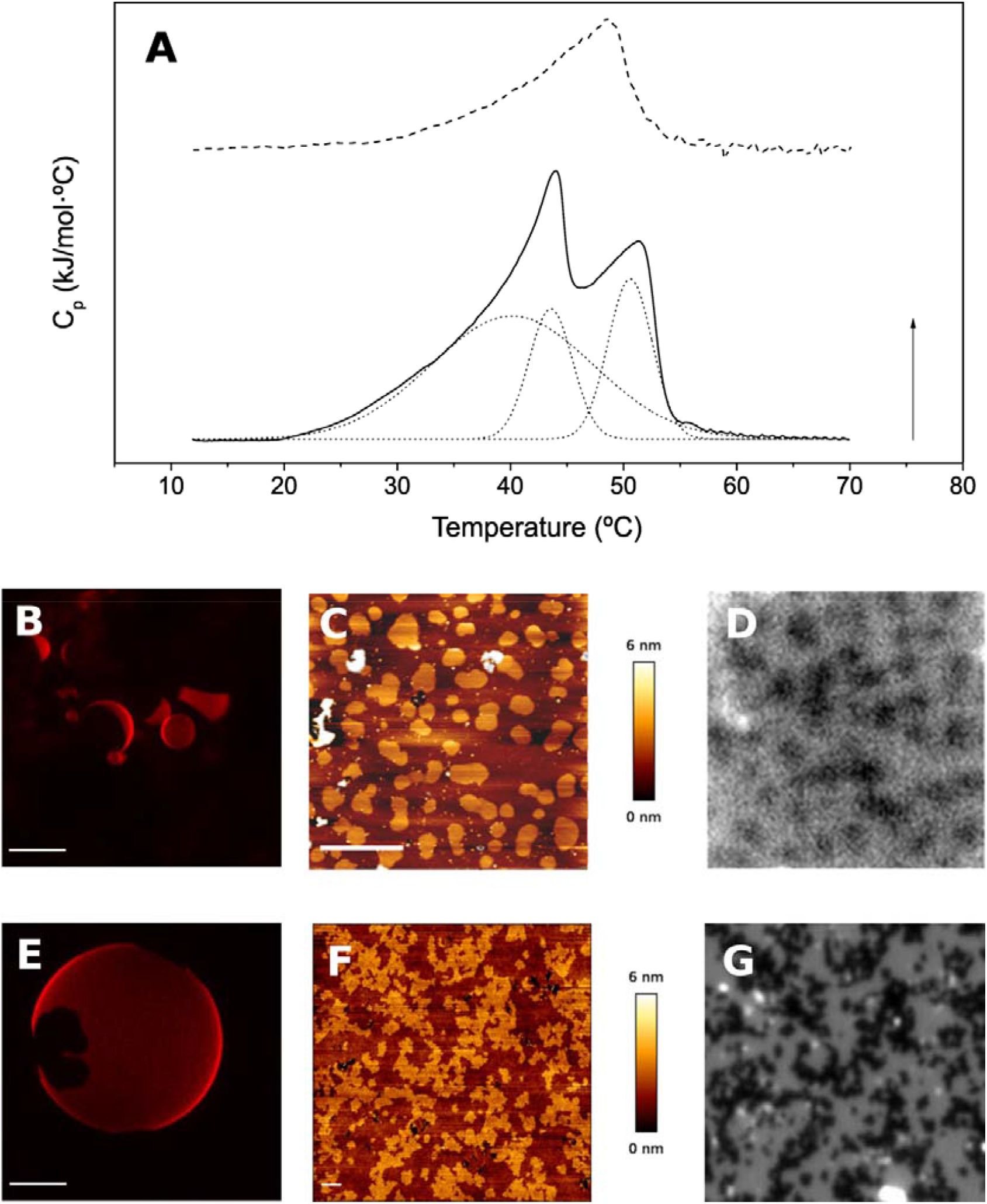
Thermograms, giant unilamellar vesicles examined by fluorescence confocal microscopy, SLB topography images obtained with atomic force microscopy, and epifluorescence images. AFM and epifluorescence images correspond to the same field. DOPC:pSM:Chol (2:1:1) + 30% pCer (A---,B,C,D) and DOPC:sSM:Chol (2:1:1) + 30% sCer (A-,E,F,G). Scale bar is 10 µm (GUV) and 5 µm (AFM). Arrow: 1.6 kJ/mol · ºC

**Table 3.** Physical characterization of DOPC:xSM:Chol (2:1:1) mixtures, with either pSM or sSM. AFM force spectroscopy: data from at least 3 independent sample preparations with at least 3 independently calibrated cantilevers. Average values ± S.D. (n=300-1000).

| Sample | x = nSM | x = pSM:nSM | x = sSM:nSM |  |
| --- | --- | --- | --- | --- |
| Phases detected in GUV | 1 | 1 | 2 |  |
| AFM breakthrough forces (nN) | | disc. $11.3 \pm 2.8$<br>cont. $5.7 \pm 1.3$ | $2.7 \pm 1.1$ | *** |

**Table 4.**
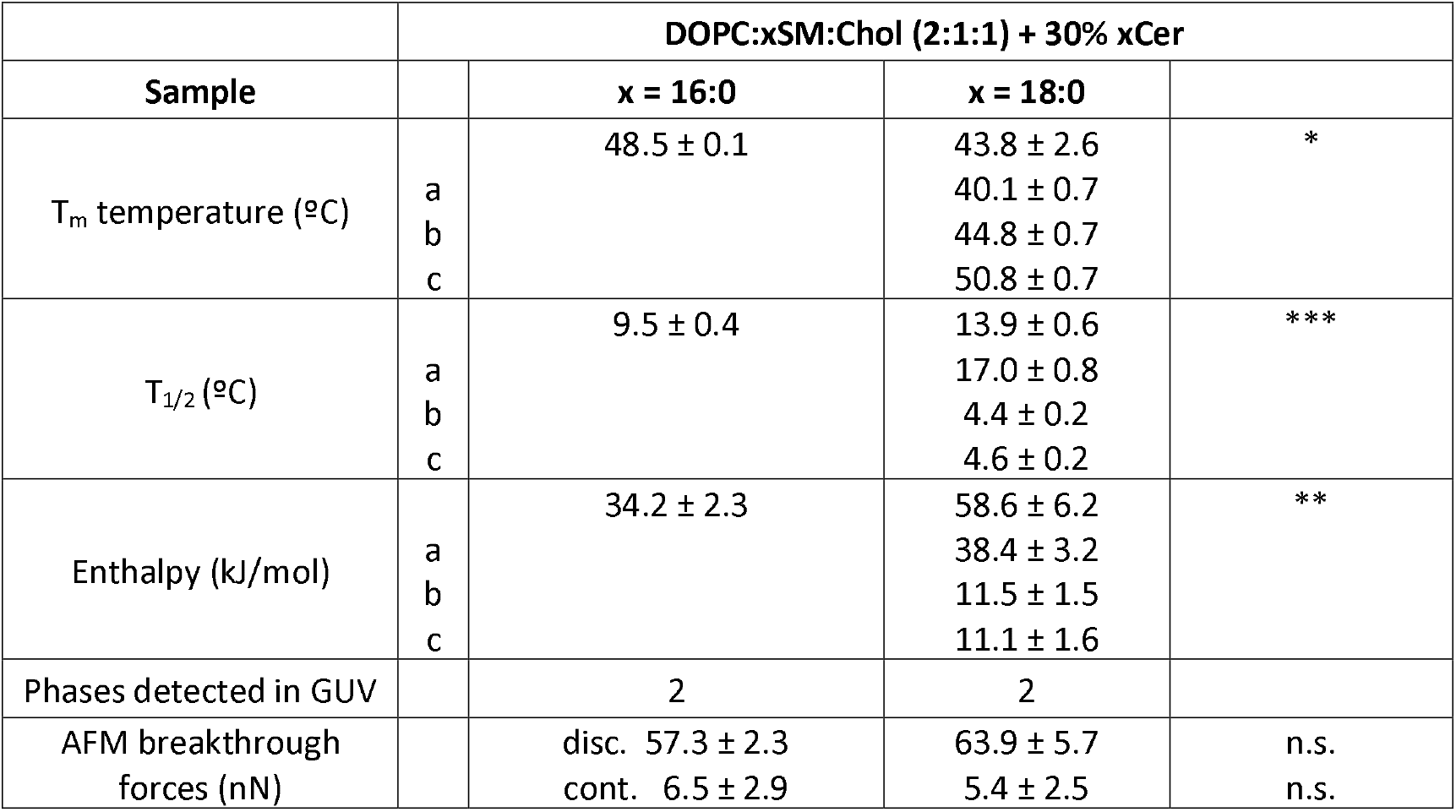
Physical characterization of DOPC:xSM:Chol:xCer (50:25:25:30) mixtures, with either C16:0 or C18:0 sphingolipid (SM and Cer). DSC data: average values ± S.D. (n = 3). The three components of the C18:0-mixture thermogram are indicated below as a, b, c. AFM force spectroscopy: data from at least 3 independent sample preparations with at least 3 independently calibrated cantilevers. Average values ± S.D. (n=300-1000).

Phase separation is detected both in the GUV (Fig. 4B, E) and in the supported bilayers (Fig.4C, D, F, G). Domains are expected to be enriched in Cer, as repeatedly observed in other samples [13,19,20,40,41]. Domain boundaries, as detected by AFM, tend to be more convoluted in the C18:0 (Fig. 4F) than in the C16:0 samples. Convoluted or abrupt/polygonal boundaries are usually taken as an indication of solid-solid phase coexistence [42]. It is not surprising that the longer-chain, saturated sphingolipids give rise to more solid phases than their shorter-chain homologs.

The epifluorescence images (Fig. 4D, G) show a good correlation with those (of the same field) provided by AFM (Fig. 4C, F). In general, the brighter, i.e. higher, domains detected by AFM correspond to darker epifluorescence areas, the latter being less accessible to the fluorescent probe. This is a general issue in all samples tested, indicating that the more rigid domains are also thicker than the more fluid ones (see color/height correlation scales in the figure). A potentially interesting difference between the C16:0 and C18:0 mixtures in these epifluorescence images is that C16:0 lipids (Fig. 4D) give rise to fuzzy dark domains, suggesting a gradient of Cer between the rigid and the fluid areas, in contrast with the sharp contours of the C18:0 domains. This in turn could be an indication of a higher Cer concentration in the fluid phase of the C16:0-based mixture, than in the C18:0 bilayers.

The AFM data provide, for the discontinuous phase, much higher breakthrough forces than for the continuous phase (Table 4). The breakthrough force for the segregated, C18:0-based domains are larger than that of those based on C16:0 lipids. The observation of higher breakthrough forces in the discontinuous than in the continuous phase is general in all the samples in which phase separation has been detected, i.e. all the 4- and 5-component ones. This is in agreement with the epifluorescence images indicating that the separated domains are more rigid, and less accessible to the fluorescent probe, than the continuous ones.

#### 3.2. DOPC:SM:Chol (2:1:1) + 30 mol% unsaturated Cer

The introduction of unsaturated SM or Cer in the 4- or 5-component mixtures (see below) makes advisable a preliminary study of a system in which only unsaturated acyl chains, i.e. no C16:0 or C18:0, are present. The DSC thermograms, GUV, AFM topography images, and epifluorescent pictures of a mixture containing DOPC:nSM:Chol (2:1:1) + 30 mol% nCer are shown in Fig. S1. In spite of the absence of saturated lipids, a clear gel-to-fluid thermotropic transition is seen (Fig. S1A), suggesting that, at variance with the situation in which saturated phospholipids, e.g. DPPC, undergo the phase transition when their acyl chains become disordered, nSM and/or nCer may exhibit a sizable endotherm when not only the acyl chains (if at all), but mainly the densely H-bonded headgroups of the sphingolipids are relaxed as an effect of temperature, and the bilayer becomes fluid. In agreement with the DSC data, the images of fluorescence microscopy and AFM reveal a clear phase separation, with discontinuous, relatively rigid domains surrounded by a continuous fluid phase (Fig. S1, B-D). Note however that, in this sample, not only does epifluorescence reveal fuzzy domain contours, but the gel domains in the GUV are less dark than in the corresponding pictures with saturated sphingolipids (e.g. Fig. 3E, 4B, E), indicating a somewhat higher fluidity of the gel phase.

In the following samples (Fig. 5), saturated SM and unsaturated Cer are combined. In this figure, pSM- and sSM-containing bilayers are compared, in both cases containing as well DOPC, Chol, and 30 mol% nCer. Their properties are summarized in Table 5.

**Figure 5.**
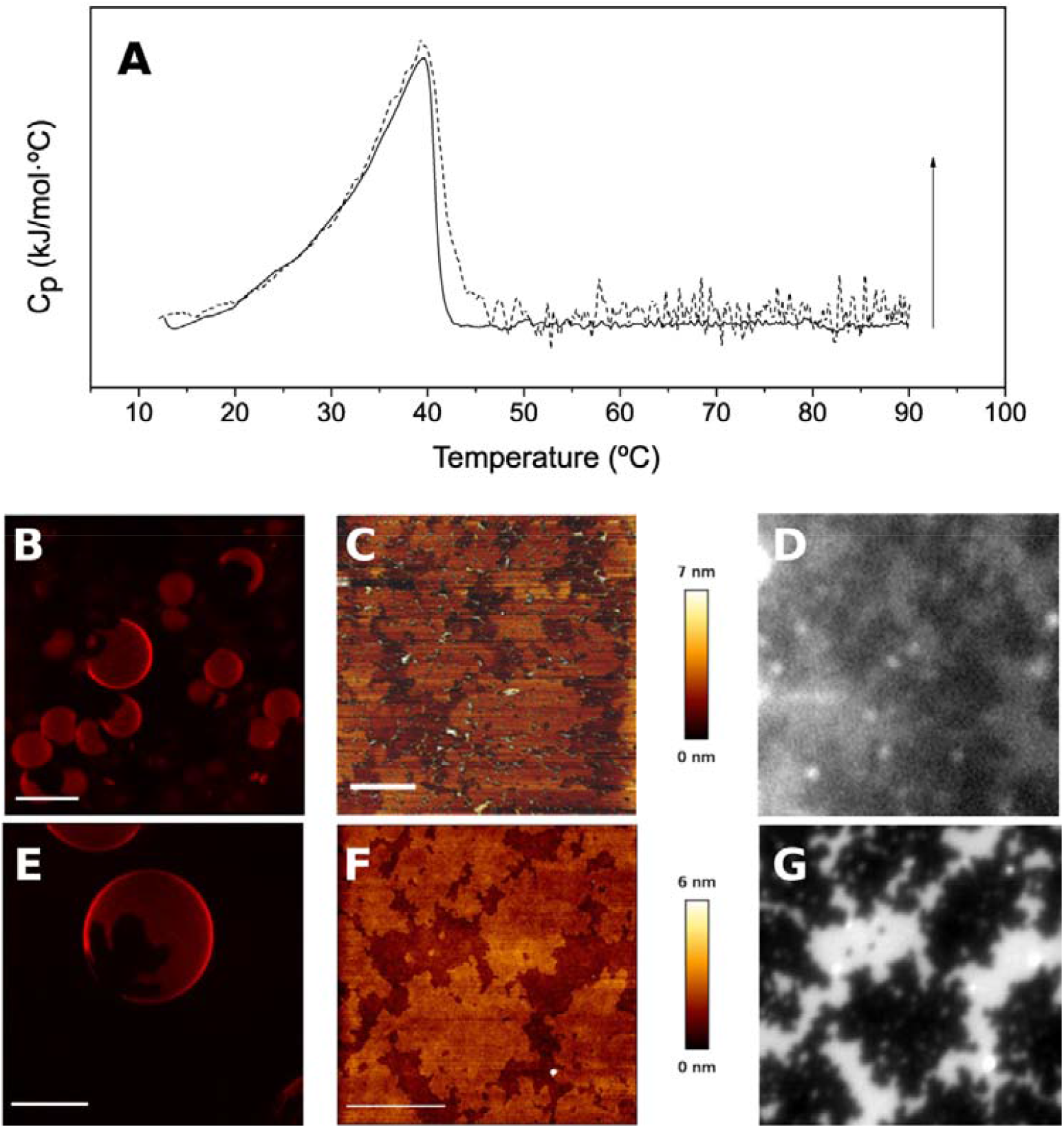
Thermograms, giant unilamellar vesicles examined by fluorescence confocal microscopy, SLB topography images obtained with atomic force microscopy, and epifluorescence images. AFM and epifluorescence images correspond to the same field. DOPC:pSM:Chol (2:1:1) + 30% nCer (A dashed line, B, C, D) and DOPC:sSM:Chol (2:1:1) + 30% nCer (A continuous line, E, F, G). Scale bar is 10 µm (GUV) and 5 µm (AFM). Arrow: 1.6 kJ/mol · ºC.

**Table 5.** Physical characterization of DOPC:SM:Chol:Cer (50:25:25:30) mixtures, with either pSM or sSM, and nCer. DSC data: average values ± S.D. (n = 3). AFM force spectroscopy: data from at least 3 independent sample preparations with at least 3 independently calibrated cantilevers. Average values ± S.D. (n=300-1000).

| **DOPC:xSM:Chol (2:1:1) + 30% nCer** |  |  |  |
| --- | --- | --- | --- |
| **Sample** | **x = 16:0** | **x = 18:0** |  |
| T<sub>m</sub> temperature (°C) | 39.0 ± 0.4 | 39.6 ± 0.1 | n.s. |
| T<sub>1/2</sub> (°C) | 8.4 ± 0.5 | 8.5 ± 0.0 | n.s. |
| Enthalpy (kJ/mol) | 24.4 ± 0.8 | 24.7 ± 3.1 | n.s. |
| Phases detected in<br>GUV | 2 | 2 |  |
| AFM breakthrough<br>forces (nN) | disc. 17.3 ± 2.0 | 17.3 ± 4.2 | n.s. |
|  | cont. 3.4 ± 1.5 | 5.8 ± 1.0 | n.s. |

Hardly any differences are seen between the pSM and the sSM mixtures in DSC or GUV data. However, in the SLB, from the point of view of AFM, the sSM continuous phase appears to be somewhat stiffer (less dark) than the corresponding pSM continuous phase (Fig. 5C, F). Also, even more than for the mixtures in Fig. 4, epifluorescence reveals for the C16:0 sample an almost continuous gradient of fluorescence (probe accessibility) between the domains and the continuous phase (Fig. 5D, G). In agreement with these observations, the breakthrough forces of the samples in Fig. 5, Table 5, are much lower (by two-thirds) than the corresponding ones in Table 4. Thus the domains in samples containing nCer are actually less rigid than in those containing C16:0 or C18:0 Cer. Otherwise, the almost overlapping thermograms (Fig. 4A) predict already the similarity of sample topologies (Fig. 4C, F). When compared with the homologous systems with saturated Cer (Table 4), both T_m_ and ΔH are decreased in the unsaturated-ceramide samples (Table 5).

#### 3.3. DOPC:xSM:Chol (2:1:1) + 30 mol% yCer

In this case, x can be either a C16:0 or a C18:0 acyl chain; y refers to either an equimolar 15% pCer + 15% nCer (Fig. 6A [discontinuous line], B-E), or to an equimolar 15% sCer + 15% nCer (Fig. 6A [continuous line], F-H). The overall mol ratios in the mixture are DOPC:SM:Chol:Cer (50:25:25:30). The physical parameters are summarized in Table 6.

**Figure 6.**
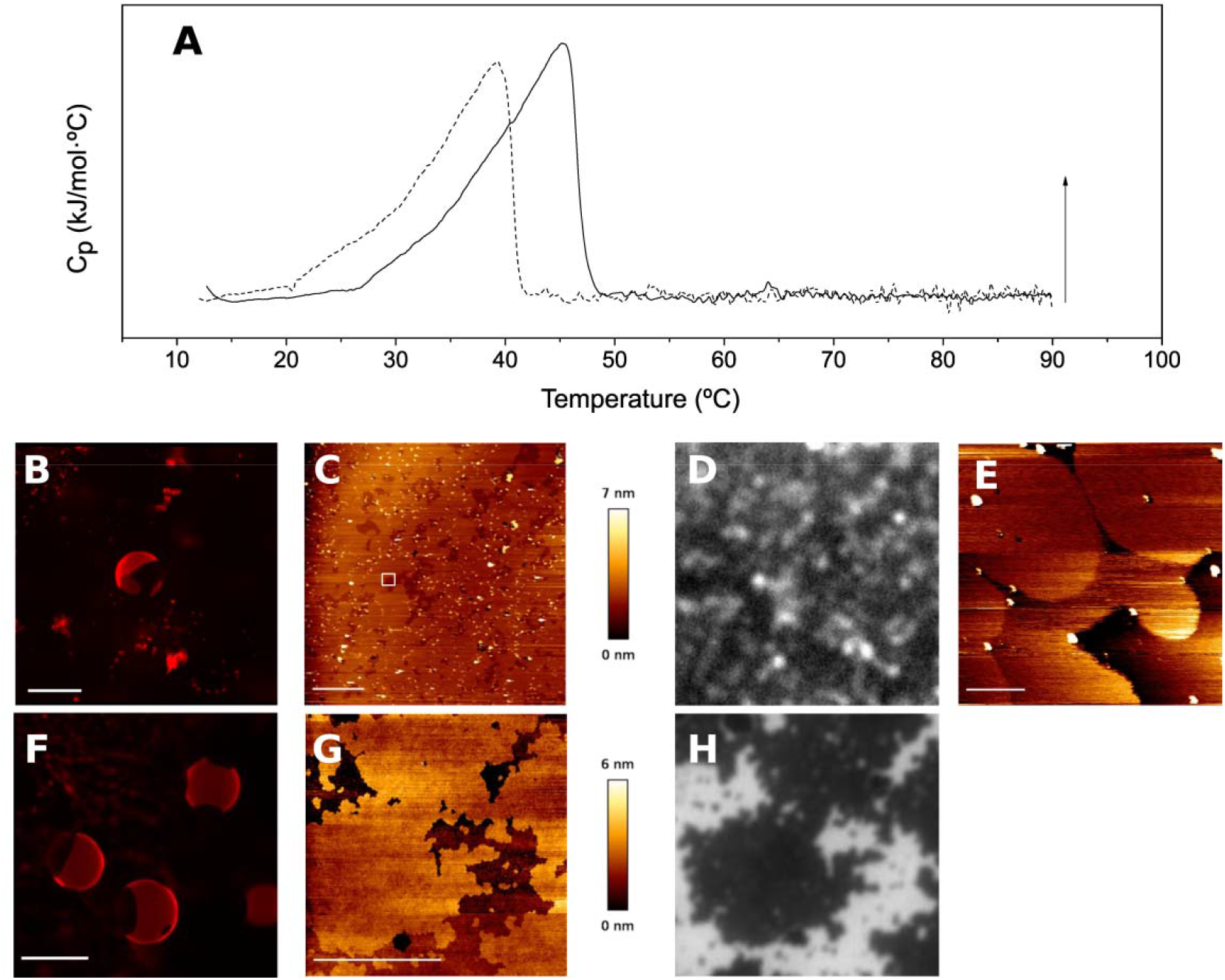
Thermograms, giant unilamellar vesicles examined by fluorescence confocal microscopy, SLB topography images obtained with atomic force microscopy, and epifluorescence images. AFM and epifluorescence images correspond to the same field. DOPC:pSM:Chol (2:1:1) + 15% pCer + 15% nCer (A dashed line, B, C, D, E), and DOPC:sSM:Chol (2:1:1) + 15% sCer + 15% nCer (A continuous line, F, G, H). Image E is a zoom-in of the highlighted area in C. Scale bar is 10 µm (GUV), 5 µm (AFM), 500 nm (AFM zoom). Arrow: 1.6 kJ/mol · ºC.

**Table 6.** Physical characterization of DOPC:SM:Chol:Cer (50:25:25:30) mixtures, with either pSM, pCer + nCer, or sSM, sCer + nCer. DSC data: average values ± S.D. (n = 3). AFM force spectroscopy: data from at least 3 independent sample preparations with at least 3 independently calibrated cantilevers. Average values ± S.D. (n=300-1000).

| DOPC:xSM:Chol (2:1:1) + 15% xCer + 15% nCer |  |  |  |
| --- | --- | --- | --- |
| Sample | x = 16:0 | x = 18:0 |  |
| $T_m$ temperature (°C) | $39.5 \pm 0.3$ | $45.2 \pm 0.1$ | *** |
| $T_{1/2}$ (°C) | $9.5 \pm 0.6$ | $9.0 \pm 0.4$ | n.s. |
| Enthalpy (kJ/mol) | $28.6 \pm 1.7$ | $27.3 \pm 2.7$ | n.s. |
| Phases detected in GUV | 3 | 2 |  |
| AFM breakthrough forces (nN) | Disc. *i* $20.9 \pm 2.5$ | Disc. $36.8 \pm 7.2$ | * |
| | Disc. *ii* $3.7 \pm 0.8$ | | |
| | Cont. $2.4 \pm 0.6$ | Cont. $3.0 \pm 1.2$ | n.s. |

When these samples are compared, the gel-fluid transition temperature of the sSM-containing mixture is higher than the corresponding pSM based bilayers, as expected from the increased chain length. An important difference is the number of phases detected in both GUV and SLB, 2 segregated phases (discontinuous i and ii) + 1 continuous phase in the pSM system, and 1 segregated + 1 continuous phase in the sSM mixtures. Fig. 5E is a zoom-in of Fig. 5C, and in it, the three phases can be distinguished. Two segregated phases had been observed in the pSM system by García-Arribas et al. (2017). It was proposed that, of the two, one was enriched in SM + Chol (L_o_-like phase), while the other one contained mainly SM + Cer (gel-like phase). The smooth, rounded areas observed at high magnification in Fig. 5E could correspond to micro- or nanodomains in an L_o_-like phase. The single segregated phase in the C18:0 samples might be due to a stronger interaction of sSM than that of pSM for the ceramides, as compared with Chol. The irregular boundaries of the sSM/sCer segregated phase (Fig. 5G, H) would be in agreement with a solid-solid lateral phase separation, and with the higher melting T of this sample (Fig. 5A).

## DISCUSSION

The results in this paper show a number of differences in the biophysical behavior of samples containing unsaturated PC, cholesterol and sphingolipids (SM, Cer), depending on whether the sphingolipid acyl chains were C16:0 or C18:0.

### C16:0 vs. C18:0

A difference of two ethylene units (‘two carbons’) in the acyl chains of a glycerophospholipid is enough to modify in a clearly detectable way its thermotropic properties, e.g. the T_m_ of DPPC and DSPC are, respectively, of 41 and 55 ºC [33]. Between 12:0 and 24:0 carbon-atom chains, the T_m_ of DPPC increases on average by 6.8 ºC per carbon atom. For the equivalent sphingophospholipid SM, the chain-length dependent increase in T_m_ per carbon atom is much lower, of 2 ºC, according to data in Jimenez-Rojo et al. (2014). This smaller dependence may be due, among other factors, to the fact that, in sphingolipids, only one of the two non-polar chains is variable, the other one corresponding to sphingosine in all cases. Moreover, in the case of SM, an increase in N-acyl length does not only lead to a higher T_m_, it also gives rise to multiple endotherms, as seen for C18:0 SM in Fig. 1B, curve I. Jimenez-Rojo et al. (2014) showed that these complex endotherms corresponded indeed to different domains, distinguishable by AFM. These authors attributed the unexpected heterogeneity in single-component bilayers to the asymmetric length of the sphingosine and fatty acyl chains, C16:0 SM being the limiting case in which the sphingoid and acyl chains would give rise to a symmetric molecule. The fact that a 16 C-atom acyl chain and a 18 C-atom sphingosine chain end up giving a molecule with two tails of similar length is due to the peculiar atomic organization of the sphingolipid polar head [43,44]. For a review on the role of H-bonding in SM interactions with other lipids in bilayers, see Slotte (2016). The polar headgroup interactions between sphingomyelin and ceramide have been recently explored with infrared spectroscopy [46]. Dupuy and Maggio (2012) studied in lipid monolayers the miscibility of ceramides with different N-acyl chains, from C10 to C18, and found that, as the hydrophobic mismatch in ceramides increased, either complete miscibility, or partial or complete immiscibility could occur depending on chain asymmetry. In a length-asymmetric molecule, the coupling of both monolayers in the membrane could be achieved either by interdigitation of the longer chains, or by curving the longer chains to accommodate them within a monolayer [48,49]. The fact that C18:0, but not C16:0, is an asymmetric molecule is probably one of the keys to interpret the results in this paper.

Table S2 summarizes the differences attributable to C16:0/C18:0 in six couples of samples (data retrieved from Tables 1-6). Examining the corresponding T_m_ (of the highest component peak when more than one was detected), it can be seen that, with one exception, compositions including C18:0 melted at higher temperatures than those containing C16:0. The ternary mixtures (not containing Cer) with C16:0 and C18:0 SM had T_m_ at 43.8 and 42.6 ºC respectively. The reasons for this behavior are not clear at present, note however that the enthalpies associated to these transitions are exceedingly small (Fig. 1, curves III and insets) and T_m_ might not be fully reliable in this particular case.

Examining the breakthrough forces of the continuous domains is interesting. For ternary mixtures (in the absence of Cer), and quaternary mixtures containing saturated Cer, the forces for penetrating the continuous phase are stronger for the C16:0 mixtures, suggesting that C16:0 SM and C16:0 Cer would be more soluble than their C18:0 counterparts in the DOPC-enriched fluid phase. The higher solubility of C16:0 sphingolipids would explain the paradox that, for these samples, the C16:0 fluid phase would be more rigid than the C18:0 one. The converse situation (C18:0 fluid phase more rigid than the C16:0 one) was found for the mixtures containing C24:1 Cer, but this ceramide is known to fluidize bilayer and counter lateral domain separation, thus leading to counterintuitive behavior of the mixtures in which it is found [40]. The discontinuous domains present a higher breakthrough force, i.e. resistance to penetration, than the continuous one, they may act as a depot of the high-melting components in the mixtures. Note also the much lower penetration forces of the fully unsaturated DOPC:nSM:Chol:nCer (50:25:25:30) mixture (Table S1), in comparison with any of the ones containing saturated sphingolipids (Tables 4-6).

The overlapping epifluorescence and AFM topography images provide additional valuable information. In general, there is a very good overlap between each couple of images (e.g. Fig. 4C and D, or F and G). However a clear, and perhaps functionally significant, difference between C16:0- and C18:0-based samples examined by epifluorescence is that, for every couple of images (Figs. 4-6, and Table S2), the domain boundaries of the C16:0 mixtures look fuzzy, while they appear clear-cut for the C18:0 homologs. It appears that the C16:0 rigid lipids (SM and/or Cer) that make up the bulk of the laterally separated domains are partly solubilized in the fluid phase, generating a sort of a gradient between both phases, while the C18:0 SM and/or Cer keep a more strict phase separation. This is particularly clear for the DOPC:xSM:Chol:xCer samples (Fig. 4 and Table 4), where a blurred domain contour is accompanied by an increased rigidity of the continuous, fluid phase when x = C16:0. In samples containing the unsaturated C24:1 Cer, that is known to counter lateral phase separation in this kind of systems (Maté), fuzzy domains are detected when C16:0 lipids are present, (Fig. 5, 6) but the corresponding fluid phases are less rigid for the C16:0 than for the C18:0 mixtures. Why epifluorescence can reveal these apparent rigidity (or lipid composition) gradients, while AFM topography detects sharp boundaries in all cases deserves further investigation.

### Physiological correlation

The above results might have implications for the physiology of the nervous system. The main sphingolipid species in brain are C18:0 SM and C24:1 SM [22,50–52][22,50,51], while in other tissues the main species are C16:0 SM and C24:1 SM [22]. Cer are present at much lower concentrations under resting conditions, about one order of magnitude, and again C18:0 Cer is the predominant species in brain, while 24-C species predominate in other tissues. The ternary mixtures described in Fig. 2 and Tables 1, 2 constitute a simplified model of the plasma membrane outer monolayer composition [53,54]. Under stress conditions, the local concentration of Cer may increase by one order of magnitude [55], the lipid composition of the outer monolayer being then represented by the quaternary mixtures in Figs. 4-6, Tables 4-6. One of the main observations in this paper is that, under most conditions, the saturated SM give rise, in conjunction with Chol and Cer, to phase-separated rigid lipid domains, coexisting with a continuous fluid phase. Of all the tested mixtures, DOPC:nSM:Chol (2:1:1) and DOPC:nSM:pSM:Chol (2:0.5:0.5:1) are the only ones not showing phase separation at the µm scale, although nanoscopic domains cannot be ruled out. The generalized presence of rigid domains in our systems must not be interpreted as implying a similar large scale phase separation in cell membranes, this being prevented by the high membrane protein concentrations, among other factors [56,57]. However, the fact that mixtures very similar to the ones naturally found in membranes lead to lateral phase separation is a very good argument supporting a similar phase separation, although perhaps at the nm scale, in cell membranes [58].

From the above data, it can be asserted that, in the nervous system, sphingolipid-rich domains will be formed with low levels of exchange with the surrounding predominant fluid phase. In consequence, the continuous fluid phase of nervous system plasma membranes will contain comparatively less SM and Cer than other plasma membranes, i.e. the thermodynamic activity of SM and Cer in these membranes will be low. SM is a rather inert molecule in itself, but Cer is metabolically very active. Apart from its role as a signal in programmed cell death [1,2], Cer greatly increases membrane permeability [5,8] thus, by containing less Cer, nervous system membranes are ensuring their stability, essential for nervous impulse generation and transmission. It is significant that, apart from nervous cells, the only other type of cell in which C18:0 sphingolipids predominate is the excitable muscle cell [59].

In conclusion, it can be stated that: (*i*) the lipids predominant in the outer monolayer of cell membranes tend to give rise to inhomogeneities, with the formation of rigid (nano)domains within a continuous fluid phase, (*ii*) inclusion of saturated Cer in the system will increase the rigidity of the segregated domains, (*iii*) C18:0-based sphingolipids, that are typical of the excitable cells, are less miscible with the fluid phase than their C16:0 counterparts, and (*iv*) the predominance of C18:0 Cer in the nervous system will contribute to the tightness of its plasma membranes, thus facilitate maintenance of the ion gradients.

## Credit authorship contribution statement

E.J.G.R. performed most of the experiments, and helped in analyzing the results. A.E performed confocal microscopy experiments. F.M.G. and A.A. had the original idea, provided funding, and wrote a first draft. All authors discussed the results and edited the manuscript.

## Acknowledgments

This work was supported in part by the Spanish Ministerio de Ciencia e Innovación (MCI), Agencia Estatal de Investigación (AEI) and Fondo Europeo de Desarrollo Regional (FEDER) (grants No. PGC2018-099857-B-I00 and No. PID2021-124461NB-I00), by the Basque Government (grants No. IT1625-22 and IT1270-19), by Fundación Ramón Areces (CIVP20A6619) by Fundación Biofísica Bizkaia, and by the Basque Excellence Research Centre (BERC) program of the Basque Government. E.J.G.R. is supported by Fundación Ramón Areces.

## Supplementary materials

**Figure S1.**
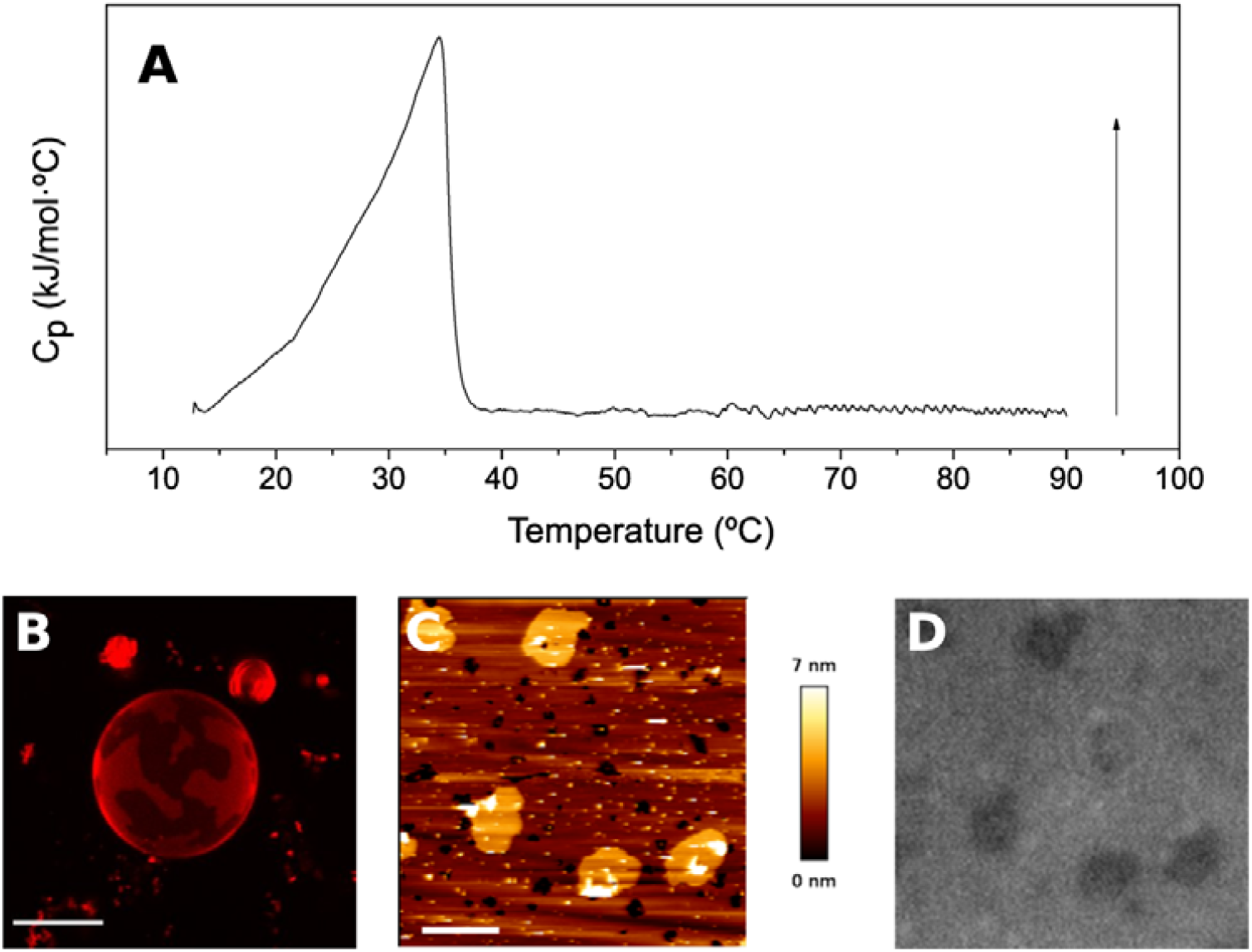
Thermograms, giant unilamellar vesicles, AFM topography images and epifluorescence images. DOPC:nSM:Chol (2:1:1) + 30% nCer. Scale bar is 10 µm (GUV) and 5 µm (AFM). Arrow: 1.6 kJ/mol · ºC.

**Table S1.** Physical characterization of DOPC:nSM:Chol:nCer (50:25:25:30) mixture. DSC data: average values ± S.D. (n = 3). AFM force spectroscopy: data from at least 3 independent sample preparations with at least 3 independently calibrated cantilevers. Average values ± S.D. (n=300-1000).

| DOPC:xSM:Chol (2:1:1) + 30% nCer |  |
| --- | --- |
| T<sub>m</sub> temperature (°C) | 31.2 ± 0.8 |
| T<sub>1/2</sub> (°C) | 8.0 ± 0.3 |
| Enthalpy (kJ/mol) | 13.0 ± 1.1 |
| Phases detected in GUV | 2 |
| AFM breakthrough forces (nN) | disc. 6.2 ± 2.4<br>cont. 1.5 ± 0.6 |

**Table S2.** A comparison of the properties of lipid compositions containing C16:0/C18:0 sphingolipids. Data taken from main text Tables 1-5.

|  | T<sub>m</sub><sup>a</sup> | Continuous<sup>b</sup> | Discontinuous<sup>c</sup> | Fuzzy<sup>d</sup> |
| --- | --- | --- | --- | --- |
| Pure SM | 41.2/44.1 |  |  |  |
| DOPC:SM (2:1) | 19.4/25.3 |  |  |  |
| DOPC:SM:Chol (2:1:1) | 43.8/42.6 | 5.7/2.7 | 11.3/2.7 |  |
| DOPC:xSM:Chol:xCer | 48.5/50.8 | 6.5/5.4 | 57.3/63.9 | ++/- |
| DOPC:xSM:Chol:nCer | 39.0/39.6 | 3.4/5.8 | 17.3/17.3 | +++/- |
| DOPC:xSM:Chol:xCer:nCer | 39.5/45.2 | 2.4/3.0 | 20.9 (3.7)/36.8 | +++/- |

<sup>a</sup> T<sub>m</sub> of the highest melting component in the sample (°C).
<sup>b</sup> AFM breakthrough force of the continuous component in SLB samples (nN).
<sup>c</sup> AFM breakthrough force of the discontinuous component in SLB samples (nN).
<sup>d</sup> Presence (+) or absence (-) of fuzzy contours in the domains seen by epifluorescence (Figs. 4-6), qualitative estimation.

